# Focal Electrical Stimulation of Human Retinal Ganglion Cells for Vision Restoration

**DOI:** 10.1101/2020.08.23.263608

**Authors:** Sasidhar S. Madugula, Alex R. Gogliettino, Moosa Zaidi, Gorish Aggarwal, Alexandra Kling, Nishal P. Shah, Ramandeep Vilkhu, Madeline R. Hays, Huy Nguyen, Victoria Fan, Eric G. Wu, Pawel Hottowy, Alexander Sher, Alan M. Litke, Ruwan A. Silva, E.J. Chichilnisky

## Abstract

Vision restoration with retinal implants that electrically stimulate retinal ganglion cells (RGCs), which transmit visual information to the brain, is limited by indiscriminate activation of many cells and cell types. Recent work in isolated macaque retina has demonstrated that direct electrical stimulation of RGCs can be performed with single-cell, single-spike resolution. However, the fidelity of epiretinal stimulation has not been examined in the human retina. Here, electrical activation of the major RGC types was examined using large-scale, multi-electrode recording and stimulation in the human retina *ex vivo* and compared directly to results from macaque. Targeted activation with single-cell, single-spike resolution was often possible without activating overlying axon bundles, at low stimulation current levels similar to those in macaque. Distinct cell types could be identified and targeted based on their distinct electrical signatures. Simulation based on these measurements revealed that a novel, dynamic stimulation approach would produce a nearly optimal evoked visual signal. These results indicate that high-fidelity control of spiking in human RGCs is achievable with extracellular stimulation and that the macaque retina is an accurate model for vision restoration with epiretinal implants.

## INTRODUCTION

Extracellular electrical stimulation of neurons has been widely used for scientific investigation of the nervous system and for the treatment of debilitating neurological conditions. Epiretinal implants, for example, electrically stimulate retinal ganglion cells (RGCs) in order to recapitulate natural visual signals and restore vision to people blinded by photoreceptor loss (*1, 2*). However, existing clinical devices provide limited restoration of visual function due to indiscriminate activation of many cells and cell types (*2*), a limitation of all existing neural implants. Recently, extensive experiments in isolated macaque monkey retina have demonstrated that RGCs can be activated precisely by direct electrical stimulation at very low current levels, in many cases with single-cell, single-spike resolution (*3–7*). These findings support the possibility of delivering high-quality artificial vision (*8, 9*), and could open the door to a new generation of high-fidelity neural implants in other parts of the nervous system. However, the high-fidelity electrical stimulation and recording demonstrated in the macaque retina has never been performed in the human retina. Thus it remains unclear whether the promising findings in the macaque can be extended to electrical activation of the human retina, and therefore whether they can be harnessed to develop high-resolution visual implants.

Here, large-scale multi-electrode stimulation and recording from hundreds of RGCs, combined with cell type identification using visual stimulation, were applied to isolated healthy human retina, and the findings were compared to results from dozens of macaque retinas in the same experimental conditions (*3, 4, 7, 8*). The results revealed electrical activation characteristics that are well-suited to high-fidelity, precise extracellular control of human RGCs, including low electrical stimulation thresholds for the major cell types, selectivity of activation, separability of cell types based on their electrical properties, and potential efficacy of real-time stimulus optimization for artificial vision. These results, which are qualitatively and quantitatively similar to findings in the macaque retina, demonstrate the ability to finely monitor and control human RGC activity, supporting the development of high-resolution epiretinal implants for treating vision loss.

## RESULTS

### Electrical and visual stimulation of identified RGC types in the human retina

To probe the focal electrical activation of the major human RGC types, simultaneous stimulation and recording with a large-scale high-density multi-electrode array system (512 electrodes, 60 μm pitch), combined with visual stimulation, were performed in an isolated human retina using procedures developed previously for macaque retina ((*3, 4, 7*), see Methods). To identify the four major RGC types in the human retina, white noise visual stimulation was performed, and reverse correlation was used to summarize the spatiotemporal receptive field properties of each cell (*10–12*). These properties, and the mosaic spatial organization of functionally identified cell types (Fig. 1, side panels), uniquely identified the ON-parasol, OFF-parasol, ON-midget and OFF-midget RGC types (*11–13*). These are the numerically dominant RGC types in the human and non-human primate retina, accounting for ~65% of the entire visual signal transmitted to the brain (*14, 15*). In this study, analysis was restricted to these four cell types.

**Figure 1.**
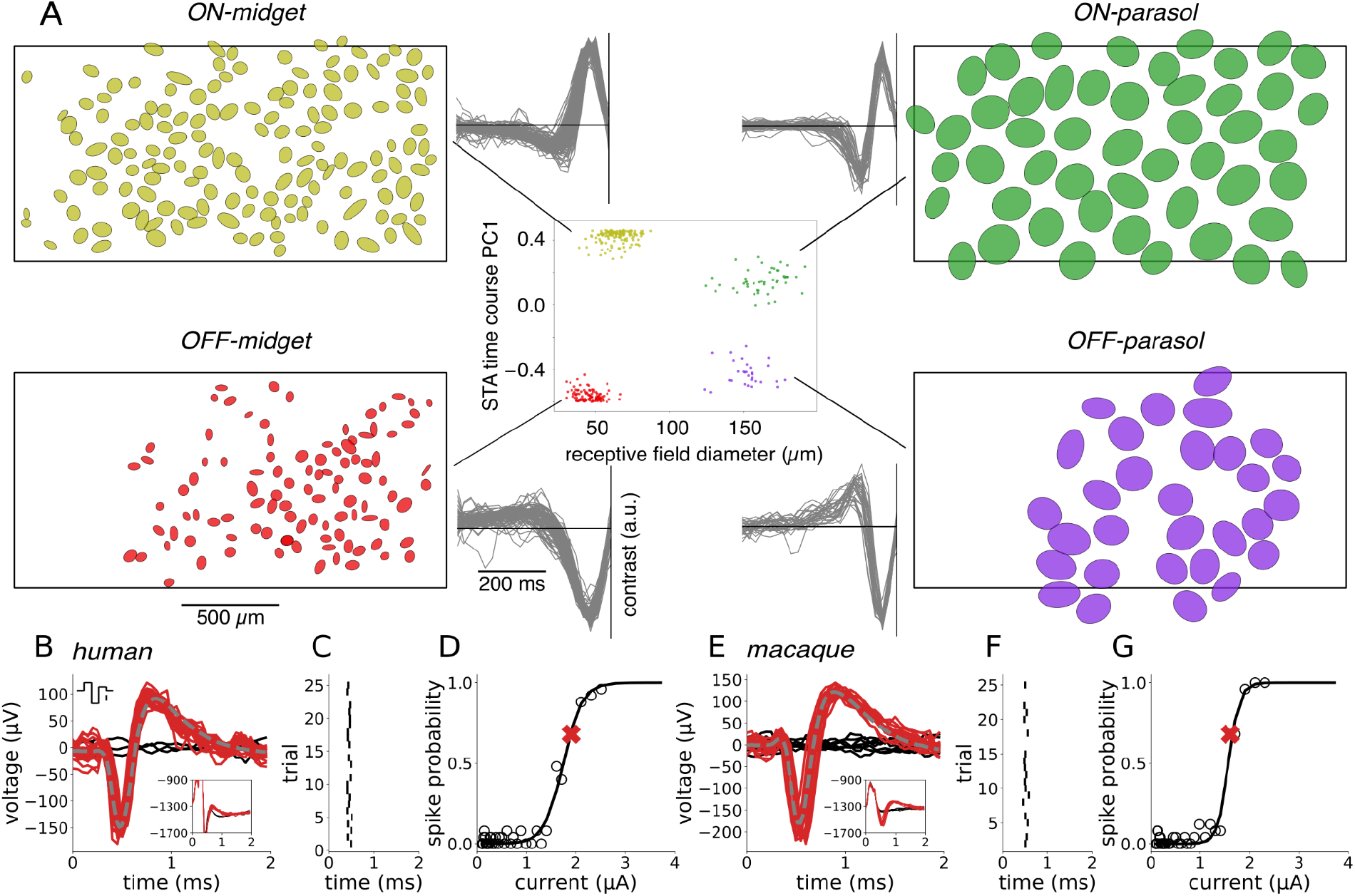
Cell type classification and spike sorting of electrical stimulation data. **(A)** Center: recorded cells were separated into distinct functional clusters by examining the receptive field diameter and the projection onto the first principal component of light response time courses obtained from the spike-triggered average (STA) stimulus. The four numerically-dominant cell types in the human retina --ON-parasol, OFF-parasol, ON-midget, and OFF-midget -- were identified from these clusters. Flanking panels: receptive field mosaics of each cell type and overlaid STA time courses of all cells from each cluster. The black rectangular outline represents the approximate location of the electrode array. **(B,E)** Artifact-subtracted voltage traces (red, black) recorded after 25 trials of stimulation with a triphasic current pulse (top inset, scale bar: 1 μA). The recorded waveforms on many trials (red) resembled the average spike waveform obtained with light stimulation (gray, dashed). Other trials resulted in no spikes (black). The artifact estimate was obtained by averaging the responses recorded in automatically identified non-spiking trials (black, inset). **(C,F)** Latency of electrically-evoked spikes (tick marks) from trial to trial. **(D,G)** Evoked spike probability as a function of current amplitude. The spike probability (open circles) was computed for each current amplitude, and a sigmoidal curve was fitted to the results (black curve, see Methods). The red “X” indicates the current level used in panels B&E.

A brief (~0.1 ms) weak (~1 μA) pulse of current delivered extracellularly through a single electrode was able to evoke a single spike in one or a few human RGCs with sub-millisecond temporal precision, raising the possibility of high-fidelity vision restoration. To quantify the probabilistic firing of a RGC in response to electrical stimulation, a triphasic current pulse (Fig. 1 B&E, top inset) was delivered repeatedly through a single electrode. In the 3 ms time window after the current pulse, the recorded voltage waveform on the stimulating electrode and nearby electrodes generally took one of two forms: electrical artifact alone, or electrical artifact with one or more evoked spikes superimposed (Fig. 1 B&E, bottom inset). These spikes were identified by clustering the recorded voltage waveforms and subtracting the average of the artifact-only traces from the traces featuring evoked responses. Each evoked spike waveform was then assigned to the cell with the most similar spike waveform recorded during visual stimulation (Fig. 1 B&E, see Methods). These steps were carried out using a novel, automated graph-based spike-sorting algorithm (see Methods). For a given cell, the variation in the time of the evoked spike from trial to trial was small (~0.03 ms SD; Fig. 1 C&F), as expected with the directly electrically elicited spikes produced in the present experimental conditions (*16*). Increases in current amplitude produced a progressively greater probability of evoking a spike on a given trial, a relationship that was summarized by a sigmoidal activation curve (Fig. 1 D&G). The activation threshold was defined as the current level that produced an interpolated spike probability of 0.5.

### Electrical activation characteristics of human RGCs are similar macaque

Spatial maps of the sensitivity to electrical stimulation for each recorded cell revealed activation at distinct cellular compartments (Fig. 2A, middle). The average spike waveforms recorded on all electrodes in the absence of electrical stimulation were used to identify the locations of somas and axons, based on their amplitude, biphasic or triphasic temporal structure, and propagation over space (Fig. 2A, top; (*17*)). These features broadly resembled the spatiotemporal spike waveforms observed in macaque RGCs (Fig. 2B, top). The spatial maps revealed that each RGC could be activated at several locations near the soma and axon (Fig. 2 A&B). In general, electrodes near the dendrites did not evoke spikes in the current range tested.

**Figure 2.**
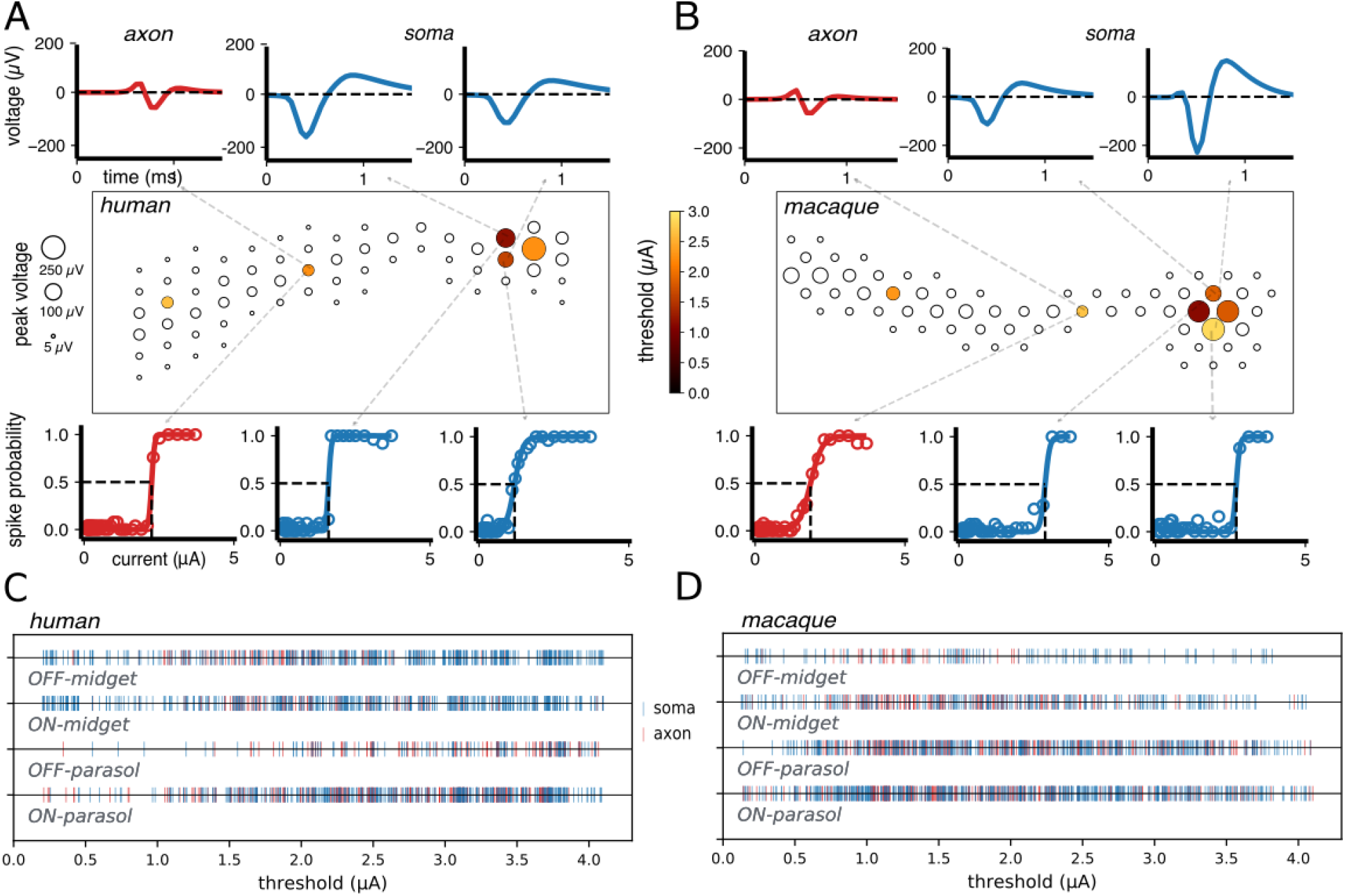
Electrical stimulation and recording of individual RGCs and populations of RGCs. **(A,B)** The electrical image (EI) of each cell is displayed in a box representing a region of the electrode array, with circles indicating the locations of electrodes on which light-evoked spikes from the cell were detected. The size of each circle represents the peak amplitude of the recorded spike at that electrode. Recorded spike waveforms are shown for three selected electrodes located near the soma (top, blue) and axon (top, red). For each cell, several electrodes (filled circles) were capable of eliciting spikes with electrical stimulation in the range of current amplitudes tested, while others were not (open circles). The colors of the filled circles correspond to activation thresholds. Activation curves are shown for each example somatic (bottom, blue) and axonal (bottom, red) recording site. The threshold response probability of 0.5 and current are indicated by dashed lines (bottom). **(C)** Activation thresholds for OFF and ON midget and OFF and ON parasol cells from five human retina recordings (1807 thresholds from 742 cells). Thresholds from stimulating electrodes located near somas and axons are indicated in blue and red, respectively. **(D)** Same as panel C, but for 18 macaque recordings (3429 thresholds from 1948 cells).

To compare electrical response characteristics across many cells of different cell types, 1807 activation thresholds resulting from somatic and axonal stimulation in the four major human RGC types were examined over 5 retinal preparations. Thresholds for all four types were in a similar range (~0.7-3.7 μA), regardless of whether the stimulating electrode was located over the soma or the axon (Fig 2 C). Activation thresholds aggregated over hundreds of RGCs from 18 macaque recordings overlapped substantially with those of human RGCs (Fig. 2 C&D; (*4*)). Specifically the mean ± standard deviation across recordings of the median threshold of each recording for OFF-midget, ON-midget, OFF-parasol, and ON-parasol cells was, respectively, 1.9 ± 0.34 μA, 2.1 ± 0.38 μA, 2.3 ± 0.43 μA, and 2.2 ± 0.44 μA across 18 macaque preparations, and 2.3 ± 0.35 μA, 2.5 ± 0.38 μA, 2.6 ± 0.49 μA, and 2.4 ± 0.35 μA across 5 human preparations.

### Human RGCs can be selectively targeted and electrically activated

To test whether it is possible for an individual human RGC to be selectively activated without activating other nearby cells, the activation curves of a collection of RGCs in a small area of a single recording were determined in response to current passed through a single electrode (Fig. 3). For each sample target cell tested, adjacent non-target cells were identified based on the proximity of their receptive fields, as a proxy for the proximity of their somas (*18*).

**Figure 3.**
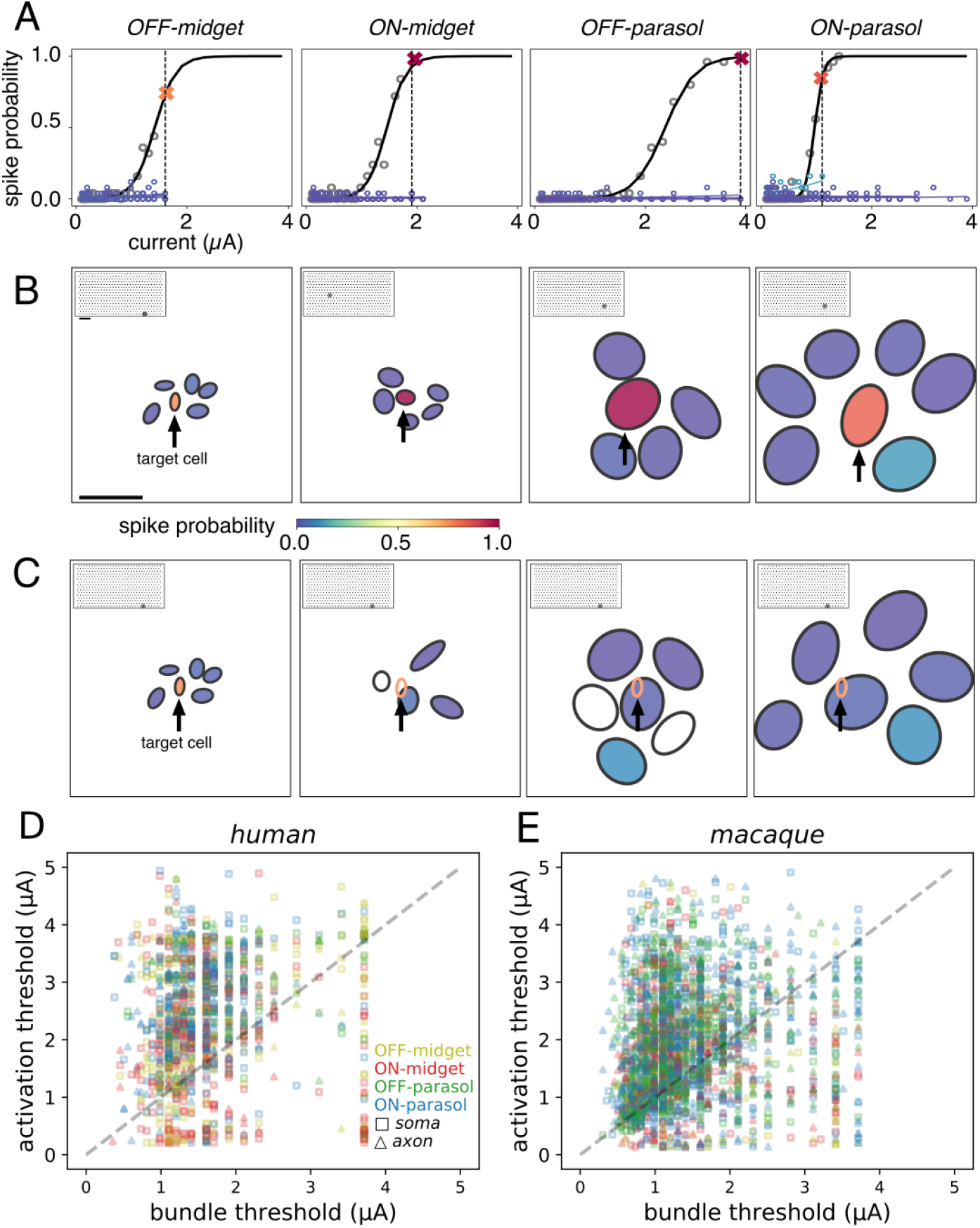
Cellular selectivity of electrical stimulation within and across cell types, and with respect to axon bundles. **(A)** Activation curves of a target cell (black) and neighboring cells of the same type (shades of blue). For each cell type, it was possible to activate a target cell (spike probability > 0.75) without activating neighboring cells of the same type (spike probability < 0.25). The largest current amplitude that produced selective activation is shown by the dashed vertical line, and the response probability of the target cell at this amplitude is marked by a red or orange “X”. **(B)** Receptive fields of the target cell (center) and neighboring cells from panel A. Receptive field fill colors indicate the spike probability at the current amplitude marked in panel A (vertical line). For each set of analyzed cells, the stimulating electrode location is represented by a gray dot on the inset array (scale bars: 250 μm). **(C)** Same as panel B, but for a single OFF-midget target cell (arrow) with neighbors of each type shown in a separate panel. In the right three panels, the target OFF-midget cell receptive field is represented by an open orange ellipse. The open black receptive fields indicate cells for which electrically elicited spikes were not identified. **(D)** Thresholds from five human retina recordings (742 total cells analyzed, 1260 cell-electrode pairs). Each electrode has a single associated bundle threshold and activation threshold, corresponding to the selectively activated cell. **(E)** Same as panel D, but for macaque retina (18 recordings, 1948 analyzed cells at 2753 distinct stimulating electrodes).

To examine selectivity *within* a cell type, the activation curve of the target cell was first compared to the activation curves of the adjacent cells in the mosaic of the same type (Fig. 3A). Note that spike sorting could only be performed reliably in a subset (742/1050) of the parasol and midget cells identified with visual stimulation (see Methods), limiting investigation to one or two regions of the retina that contained locally complete mosaics of each cell type. Examples of target cell activation at current amplitudes that did not activate any of the neighboring non-target RGCs of the same type were identified for each of the four cell types (Fig. 3B), and represented the pattern observed in the overwhelming fraction (~98%) of activated cells of each type.

To further test whether selectivity *across* cell types could be achieved, an example target OFF-midget cell and RGCs of all four major types with overlapping or nearby receptive fields were analyzed. Selective activation of the target OFF-midget cell was achieved without activating nearby cells of the other major types (Fig. 3C), similar to previous findings in the macaque (*4*). To determine the prevalence of this kind of selective activation, activation thresholds of all measured cells were collected across cell types and preparations. Out of the 22%, 24%, 42%, and 41% of all recorded OFF-midget, ON-midget, OFF-parasol, and ON-parasol cells that could be activated using at least one electrode delivering current within the tested stimulus amplitude range, 71%, 81%, 93%, and 89% respectively could be activated without eliciting spikes any other recorded cell.

Reconstituting high-resolution vision requires not only selective activation of individual RGCs in local regions, but also potentially requires avoiding axons that run in bundles across the retinal surface and convey visual signals from distant, unidentified cells [9]. To test whether this is possible, bundle activation thresholds were measured at each electrode. This was accomplished by identifying the lowest current amplitude that produced bi-directionally propagating electrical activity (*7*) that reached opposite edges of the array (rather than terminating at a soma on the array) (*19*). Bundle activation thresholds were then compared to RGC activation thresholds on the same electrodes. Most (73%) electrodes activated bundles at current levels below the activation thresholds of nearby RGCs (Fig. 3A), confirming the substantial challenge of bundle activation (see Discussion).

To determine the potential impact of axon bundles on the ability to selectively activate single cells, activation thresholds were compared to bundle thresholds. In human retina preparations, 228/742 (31%) cells (all four types) were selectively activated below the bundle threshold at 279/1260 stimulation electrodes tested. Comparison to data from 18 macaque preparations revealed a similar distribution of RGC and bundle activation thresholds (Fig. 3B), with 664/1948 (34%) cells selectively activated at 869/2753 stimulation electrodes without bundle activation. In summary, although undesirable axon bundle activation limits the selectivity of epiretinal stimulation, with appropriate calibration, the electrodes that activate individual RGCs at current levels below bundle threshold can be identified and used (see Discussion).

### Identification of cell types for optimal electrical stimulus delivery

Effectively utilizing high-resolution electrical stimulation in a clinical implant requires identifying and targeting functionally distinct RGC types without the ability to measure light-evoked responses. In the macaque retina, cell type identification using only electrical properties has been demonstrated (*6*). To test whether this is also possible in the human retina, electrical features were used to distinguish the four major RGC types. Axon conduction velocity, estimated from the EIs of recorded cells (Fig. 4A, see Methods), correctly distinguished 250/251 parasol and midget cells in one recording (Fig. 4A), and 141/149 cells in a second recording (not shown) (*20*). Within the parasol and midget cell classes, the autocorrelation function (ACF) of the recorded spike train was used to distinguish ON and OFF cells (*12, 21, 22*). The ACFs of ON-parasol and OFF-parasol cells were visibly distinct, and *k*-means clustering (*k* = 2) of projections onto the first seven principal components enabled cell type separation that was correct for 86/89 cells in one recording (Fig. 4B) and 72/84 cells in a second recording (not shown). Discrimination of ON-midget and OFF-midget cells based on ACFs was ineffective in one recording (277/423 cells, not shown), but was effective in a second recording (176/184 cells, Fig. 4C). Although this variability requires further characterization and may pose a challenge in clinical implants (see Discussion), the results demonstrate that recorded electrical features of cells, such as EIs and ACFs, have the potential to enable cell type identification.

**Figure 4.**
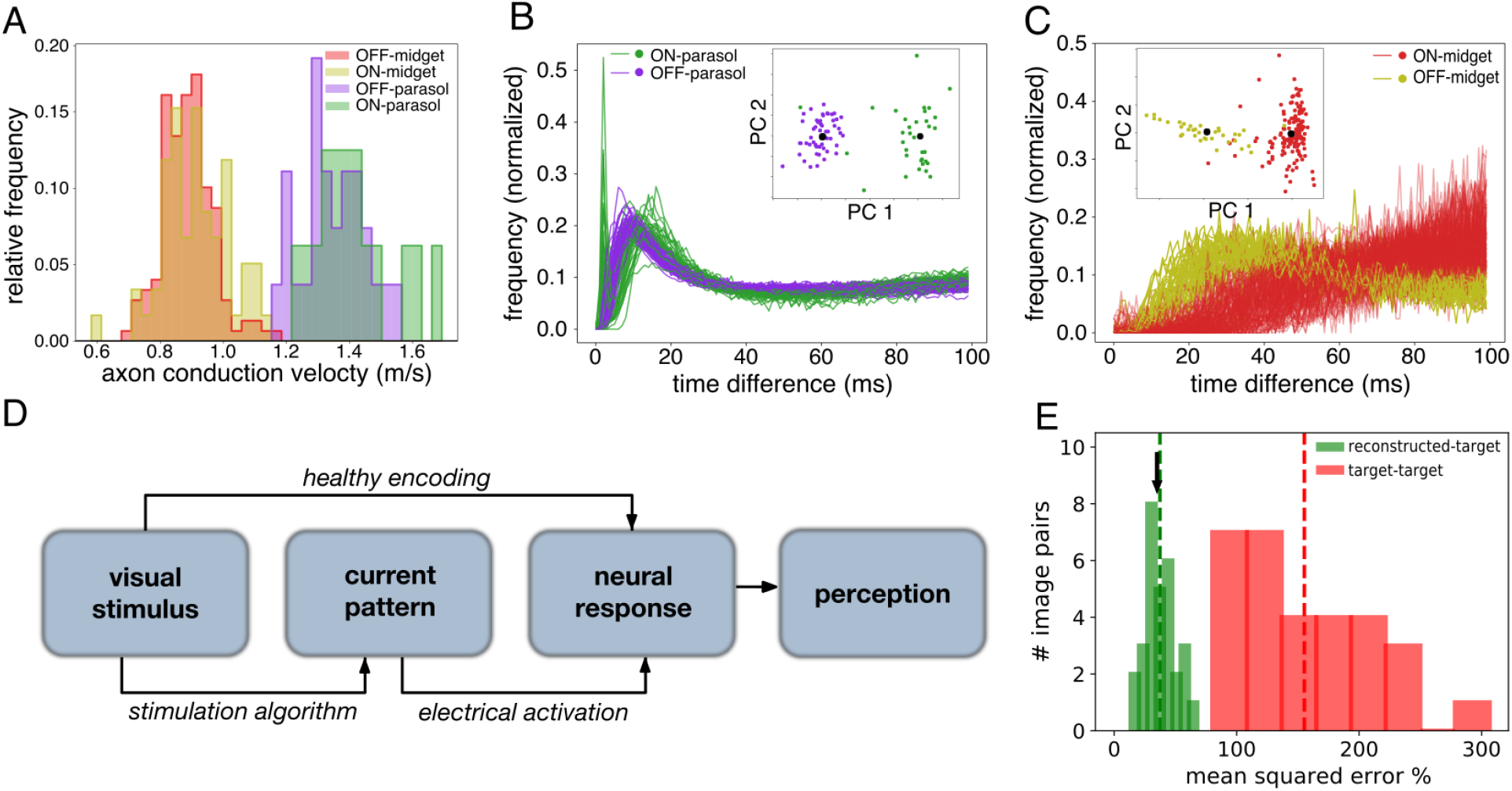
Identification of the four major RGC types and performance of a simulated stimulation procedure. **(A)** Histograms of axon conduction velocities for 251 ON-parasol, OFF-parasol, ON-midget and OFF-midget cells computed from electrical images. **(B)** Spike train autocorrelation functions for 89 ON and OFF parasol cells. Inset: clustering of projections onto the top 7 principal components separates ON from OFF cells. Black circles represent centroids of each cluster. **(C)** Same as panel B, but for 184 ON and OFF midget cells. The same recording was used for panels A and B, but a separate recording was used for panel C. **(D)** Schematic representation of a perceptual image reconstruction paradigm to guide prosthetic stimulation (see Results). **(E)** Relative mean-squared error of predicted reconstruction using a temporally dithered stimulation algorithm with respect to 15 randomly generated target images (green bars; dashed green line shows mean), compared to the error between randomly chosen target images (red bars and line), and the mean error of optimally chosen stimulation sequences (blue arrow).

To assess the potential application of the above findings to vision restoration, a newly developed stimulation algorithm and assessment metric were used (Fig. 4D, (*9*)). To produce high-fidelity artificial vision, precise RGC activity evoked by individual electrodes must be combined across the array to reproduce the rich spatiotemporal response patterns evoked by natural visual stimuli. However, in general this cannot be achieved simply by passing current through multiple electrodes simultaneously, because currents delivered by different electrodes can combine nonlinearly to influence firing (*5*). A recent approach uses *temporally dithered* current pulses (*9*) to address this problem, exploiting the relatively slow temporal integration of visual perception (tens of milliseconds, see (*23*)) compared to the high temporal precision of epiretinal stimulation (~0.1 ms) (Fig. 1 C&F). Briefly, a target visual stimulus is converted into a sequence of single-electrode stimuli, each probabilistically evoking spikes in one or more target cells. The perceived reconstructed image is assumed to depend only on the total number of spikes generated by each cell within the integration time of the visual system (here, we assume 50 ms). This approach was simulated using measured activation curves and a linear decoding model of the resulting perceived image, with a sequence of stimuli chosen to reduce the error maximally at each time step. This procedure resulted in an expected mean-squared error of 37% relative to the random checkerboard target images (Fig. 4E, green bars and line). This was substantially smaller than the error between two randomly selected target images (170%; Fig. 4E, red bars and line) and only slightly larger than a lower bound on the optimal algorithm (36%; Fig. 4E, blue arrow; see Methods for details). The similarity of these results to previous findings in macaque retina (*9*) suggests that this approach may translate effectively for vision restoration in humans.

## DISCUSSION

The present results reveal that high-fidelity electrical stimulation of RGCs in the human retina is possible, a potentially crucial finding for epiretinal implant design – closely matching findings in macaque retina. Electrical activation of the four numerically dominant cell types -- ON-parasol, OFF-parasol, ON-midget and OFF-midget -- with sub-millisecond precision and low activation thresholds, was very similar to macaque (Fig. 1, 4; (*4*)). In some cases, activation of a single RGC without activating neighboring RGCs (Fig. 3; (*4*)) or axon bundles (Fig. 3; (*7*)) was possible, providing evidence for cellular-level control of the neural code. It was also possible to distinguish the four major cell types using only electrical features (Fig. 4; (*6*)) rather than light response properties which would not be available in a blind person. Finally, simulation of a temporal dithering strategy approached the performance of an optimal stimulation algorithm (*9*). Below, we discuss the applicability of the present results to a future retinal implant, as well as caveats and potential extensions.

The results suggest that future retinal implants could exploit focal RGC stimulation to provide high-fidelity artificial vision to the blind. Existing implants that directly stimulate RGCs are capable of eliciting coarse light percepts, but provide little restoration of useful vision (*1, 2, 24*). A major limitation of these devices is that their large stimulating electrodes result in indiscriminate activation of multiple cells and cell types. A faithful replication of the neural code of the retina instead requires the ability to stimulate individual RGCs of each major type in distinct, naturalistic temporal patterns. The findings in both the macaque retina (*3, 4, 7*) and in the human retina collectively demonstrate the possibility of such control over RGC activation, in some cases with single-cell, single-spike precision.

Axon bundles pose a major challenge for accurately reproducing the neural code with epiretinal stimulation: only a minority of targeted RGCs (Fig. 3A) could be activated without activating axon bundles, as in macaque retina (Fig. 3B), and consistent with perceptual reports from people with existing retinal implants (*25*). Previous work in macaque retina showed that nearly half of the cells that could be activated without activating their neighbors could also be activated without activating axon bundles (*7*). Although this proportion was difficult to estimate reliably in the human data because of the limited number of analyzable cells, the present findings are qualitatively consistent with the macaque findings, because cells with activation thresholds lower than those of their neighbors also tend to have thresholds lower than those of axon bundles. As in the macaque data, the findings suggest that many electrodes will not be able to avoid activating axon bundles, and therefore that a calibration procedure to identify and avoid those electrodes will be crucial for a clinical implant. An alternative approach could involve surgically severing incoming axon bundles on the distal side of the array during device implantation, inducing axon degeneration. Another approach is to implant the device at other retinal locations such as the raphe region, which has a relatively low density of axon bundles This may be important for targeting the human retina: results in macaque indicate that a larger fraction of RGCs in the raphe can be activated without axon bundles (*7*).

High-fidelity visual reconstruction relies on cell type identification, but the light response properties that can be used for this purpose in a healthy retina (Fig. 1; (*13*)) are unavailable in the retina of a blind person. Previous work in macaque retina (*6*) and the present work (Fig. 4) show that features of the electrical image (EI) and the spike train autocorrelation function (ACF) can be used to distinguish the four numerically dominant cell types, without relying on light responses. One feature of the EI -- axon conduction velocity -- reliably separated parasol cells from midget cells (Fig. 4A) consistent with well-known biophysical distinctions between these cell types in the macaque retina (*20, 26*). Features of the ACF separated the ON and OFF types within the parasol and midget cell classes (Fig. 4 B&C). However, because the temporal structure of ACFs varies between retinas, these features may be difficult to use for identifying (rather than merely distinguishing) cell types (see (*12, 21*)). This suggests that further analysis of EI features, or patient feedback about the perceived brightness of phosphenes evoked by electrical stimulation, could be important for identifying cell types in the clinic. In addition, ON and OFF midget cells were reliably separated based on ACFs in one recording but not in the other (see Results), again motivating further investigation of EI features. Finally, cells of other types were not considered in this analysis, but may be important for the function of a clinical implant.

Changes in the retina during degeneration could introduce additional challenges. First, the electrical activity used here for cell type identification was light-driven, whereas an implant in a blind person would rely on recording spontaneous activity. Spontaneous activity in the major RGC types in anesthetized macaques is high (20-30 Hz) (*27*), but has not been measured in humans. Some studies have indicated that spontaneous RGC activity increases during retinal degeneration (*28–30*), however, results in a rat model of photoreceptor degeneration indicate that spontaneous activity increases in OFF cells but declines in ON cells (*31*), Second, it is unclear whether the structure of the ACF changes during degeneration. It seems unlikely that ON and OFF cell ACFs would become much more similar to one another, unless the distinctions between them in the healthy retina are primarily a result of presynaptic activity that is lost during degeneration, which is unlikely based on previous work (*31*). However, changes during degeneration could make it difficult to identify (as opposed to distinguish) cell types based on ACFs. Third, it is unclear how EIs would be affected by degeneration, although their overall form probably depends mostly on the morphological structure and ion channel distribution of the cell, which may remain relatively stable (but see (*32, 33*)). Finally, for clinical application it will be critical to verify that electrical stimulation thresholds in the degenerated retina are comparable to those in the healthy retina, as has been previously demonstrated in a rat model of retinal degeneration (*34*).

Caveats about the extent of the present data are also worth highlighting. Due to the scarcity of healthy human donor retinas, analysis of electrical responses was restricted to only five human retinal preparations from three donors. Although comparison to the macaque revealed many similarities, the data are insufficient to quantify the variability across human retinas.

The present results are compatible with combining the activation of individual RGCs over space and time to reproduce the complex spatiotemporal patterns of activity that occur in natural vision. Prior work in macaque has shown that spatiotemporal patterns of activity in small groups of cells can be reproduced with high fidelity (*8*). More recently, closed-loop methods have been developed to minimize the difference between a target visual image and an estimate of the perceived image obtained through electrical activation, based on pre-calibrating the activation of RGCs of different types and temporal dithering during stimulation (*9*). The performance of this method in the present human data resembles results in macaque (see Results), suggesting that it can be applied to a future implant.

We speculate that, given the putative biophysical similarity between RGCs and spiking neurons in the brain (*35, 36*), the two principal results of this study – fine extracellular control of spiking in human neurons and quantitative similarity to the macaque – will extend to other parts of the nervous system. This could significantly advance our understanding of human neural processes and speed the development of clinical implants to treat a wide range of nervous system disorders.

## MATERIALS AND METHODS

### Retinas and preparation

Human eyes were obtained from three brain-dead donors (29 year-old hispanic male, 27 year-old hispanic male, 47 year-old caucasian female) through Donor Network West (San Ramon, CA). Macaque eyes were obtained from terminally anesthetized animals sacrificed by other laboratories, in accordance with Institutional Animal Care and Use Committee guidelines. Following enucleation the eyes were hemisected, the vitreous was removed in room light, and the eye cup was stored in oxygenated Ames’ solution (Sigma) at 33°C. Electrophysiological data were recorded from portions of the retina, as described previously (*4, 7*). Small (~3×3mm) segments of peripheral retina (10-14 mm temporal equivalent eccentricity from the fovea) were isolated from the sclera, detached from the pigment epithelium, and held RGC side down on a multielectrode array (see below). In one recording (Fig. 4 A&B) the pigment epithelium remained attached (12 mm from the fovea on the superior vertical meridian). For macaque preparations, the retina was detached from the pigment epithelium and eccentricities ranged from 8 to 12 mm temporal equivalent (*11*).

### Electrophysiological recording and visual stimulation

Recording and stimulation were performed with a custom 512-electrode (10μm diameter, 60μm pitch, 1.7mm^2^) recording and stimulation system sampling at 20kHz, as described previously (*7, 37–39*). Spikes recorded from individual neurons during visual stimulation were identified and segregated using standard spike sorting techniques (*37, 40, 41*). To identify the type of each recorded cell, the retina was visually stimulated with a binary white noise stimulus, and the spike-triggered average (STA) stimulus was computed for each RGC (*10, 11*). The STA summarizes the spatial, temporal, and chromatic structure of the light response. The spatial receptive fields and time courses obtained from the STA were used to identify the distinct cell types (Fig. 1), as described previously (*11–13, 21*). Electrical images (EIs), obtained by averaging thousands of spatiotemporal voltage patterns over the array at the times of the identified spikes from each RGC (*17, 37*) were used to infer the spatial location of the soma and axon (Fig. 3) and to identify electrically elicited spikes (see below) (*4*).

### Electrical stimulation and spike sorting

Electrical stimuli were triphasic current waveforms, consisting of a negative current stimulating phase lasting 50 μs (one sample) preceded and followed by charge-balancing positive phases of equal duration, with relative amplitudes 2:-3:1 (Fig. 1 A&D, top inset), passed through one electrode at a time. At each electrode, forty current amplitudes in the range 0.1 to 4.1 μA were delivered twenty-five times each (*4, 7*).

An automated spike sorting method was used to identify human and macaque RGC responses to electrical stimulation. The method resembles the procedure used to classify spikes manually in previous work (*3–5, 7, 8, 16*). Briefly, for each stimulating electrode, a set of potentially activatable target RGCs was identified, based on light-evoked spike waveforms featuring negative peak amplitudes larger than the noise threshold for that electrode (twice the standard deviation of the recorded electrical noise, see below). For each target RGC, a set of recording electrodes with light-evoked spike waveforms exceeding the noise threshold was identified. Then, for each stimulus amplitude, across all recording electrodes for all target RGCs, voltage traces recorded in the 3 ms period following current pulses delivered by the stimulating electrode were collected and grouped using unsupervised clustering. For each pair of clusters, signals were subtracted, aligned, and averaged to obtain residuals. These residuals were iteratively compared to each of the recorded light-evoked waveforms of all target cells on all recording electrodes to identify the set of cells contributing to each stimulation trial for each amplitude. The clustering approach enabled the algorithm to identify spikes in recorded traces containing bundle activity, because the latter is consistent across stimulation trials [9] and therefore was identified as an electrical artifact. The accuracy of the algorithm was assessed by comparing identified activation curves with activation curves obtained using manual spike sorting [6,9]. Across 227 activation curves (see below) obtained from 5 different human retina preparations and 1657 activation curves across 18 different macaque retina preparations, 94.3% of automatically identified thresholds were within 0.2 μA of the corresponding manually identified threshold.

To enable accurate spike assignment, only cells with recorded absolute spike amplitudes of at least 60 μV on at least one electrode were analyzed, because spikes with lower amplitudes were difficult to visually distinguish from noise. This threshold was empirically supported by quantification of the automated spike sorting error rate (relative to manual spike sorting) as a function of recorded spike amplitude. Even after filtering cells by this threshold, in some cases large electrical stimulation artifacts, axon bundle activation, or other biological activity made it difficult to assign spikes to identified cells with confidence. These cells were manually examined and excluded from further analysis. Of the 1050 parasol and midget cells identified during visual stimulation across human retinal preparations, 742 were analyzable by the above criteria.

For each analyzed cell, spike probabilities were computed over the twenty-five repeats of each stimulus, and were modeled as a function of current amplitude by a sigmoidal relationship:

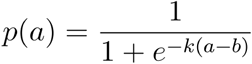

where *a* is the current amplitude, *P*(*a*) is the spike probability, and *k* and *b* are free parameters. Fitted curves were used to compute the activation threshold, defined as the current amplitude producing a spike probability of 0.5,.

### Estimation of axon bundle activation thresholds

For each stimulating electrode, axon bundle activation thresholds were estimated using a method previously described (*19*). Bundle activation can be detected by bidirectional propagation of an evoked electrical signal that, unlike the activity of a single cell, increases progressively in amplitude with increasing stimulation current (*7*). Human observers visually inspected a movie of recorded voltage on the array for a period of 2 ms after electrical stimulation. Bundle activation threshold was defined as the lowest stimulus amplitude at which bidirectional activity that increased with stimulating current reached at least two edges of the array, implying that the signal originated at least partly in the axons of cells with somas off the array. For a small number of electrodes near the edges of the array, bidirectional activity was not easily discerned. At these electrodes, the bundle activation threshold was defined as the lowest current level at which unidirectional activity grew in amplitude with current level, was clearly larger than the typical single-axon signal obtained from EIs (~35 μV), and propagated to a single distant edge of the array.

### Distinguishing midget and parasol cells using axon conduction velocity

The axon conduction velocity of each cell was estimated using axonal EI waveforms upsampled tenfold in time to improve peak spike time estimation (Fig. 3, top; see (*17, 20*)). Among all the electrodes recording axonal signals from a given cell, the distance between each pair of electrodes was divided by the corresponding time difference of the negative peaks of the EI waveforms, producing a collection of velocity estimates. The conduction velocity for the cell was estimated by computing a weighted average of these pairwise estimates, with the weight for each pair of electrodes given by the product of the peak EI amplitudes on the electrodes. Electrode pairs were excluded if the difference in time at minimum voltage was less than the sampling interval (0.05 ms). Only velocity estimates corresponding to the largest ten weight values for each cell were used. EIs that were visibly corrupted by errors in spike sorting or that contained fewer than eight axonal electrodes were excluded from analysis. With this filtering, analysis was restricted to 251 of the 520 cells in one recording and 149 of the 270 cells in a second recording.

### Distinguishing ON and OFF cells using ACFs

The autocorrelation function (ACF) of the recorded spike times for each cell was used to discriminate ON and OFF cells (see (*12, 21*)). For each cell class (midget, parasol) principal components analysis was performed on the L2-normalized ACFs of all cells, and *k*-means clustering (*k* = 2) was performed on the projections onto the top seven principal components. A cell was considered correctly separated if it was assigned to a cluster in which its type was the most numerous.

### Analysis of temporal dithering for optimizing artificial vision

The temporal dithering algorithm uses the measured responses to electrical stimulation to select a sequence of stimuli intended to produce perception matching a target visual image as closely as possible (*9*). This approach is based on the assumption that visual perception is determined by the total number of spikes from each RGC within the integration time of the visual system (~50 ms; see (*23*)), during which many stimuli can be applied. Specifically, the approach requires a *dictionary* containing elements representing the spike probabilities of all cells for each distinct electrical stimulus tested (in this case, each electrode and current level), a target visual image, and a decoder which computes an estimate of the visual image from the spike counts of all cells during the integration time window. Responses of ON and OFF parasol cells that could be stimulated by at least one electrode were used to construct the dictionary. Due to the limited number of analyzable cells, dictionary elements that activated axon bundles were permitted in the analysis, although this is not appropriate for a clinical implant. Random checkerboard images were chosen as targets for reconstruction, and the scaled spatial receptive field of each cell was chosen as the linear decoding filter for its spikes (see (*42*)). To simulate the temporal dithering approach, a sequence of electrodes and stimulating current amplitudes was chosen greedily, maximizing the expected reduction in mean squared error between the target and linearly decoded image in each stimulation time step. The performance of the algorithm was measured as relative mean squared error (mean squared error divided by mean squared intensity of the target image) over the pixels covered by the receptive fields of all cells used. The error of the algorithm was compared to a lower bound on the error of an optimal algorithm, in which the entire stimulation sequence was optimized to minimize the error between the target and decoded image (*9*).

## Contributions

SM and AK performed dissections and collected the data, SM, ARG and EJC conceived and designed the experiments, SM, ARG, RV, MZ, GA, NPS, MH, VF, HN, EGW, and EJC analyzed the data, PH, AS and AML developed and supported recording hardware and software, RAS performed the surgery, EJC supervised the project. EJC, SM, ARG, MZ, NPS, GA, and RV wrote the paper.

## Acknowledgements

Human eyes were provided by Donor Network West (San Ramon, CA). We are thankful for the cooperation of Donor Network West and all of the organ and tissue donors and their families, for giving the gift of life and the gift of knowledge, by their generous donations. We thank J. Carmena, T. Moore, W. Newsome, M. Taffe, S. Morairty, J. Horton, and the California National Primate Research Center for providing macaque retinas. We thank K. Jensen, M. Allen, and K. Tinajero, K. Berry, K. Williams, B. Morsey, J. Frohlich, and M. Kitano for helping obtain access to macaque retinas and metadata. We thank D. Palanker, F. Rieke, S. Mitra, P. Li, T. Carnevale, M. Hines, F. Rattay, P. Tandon, E. Wu, R. Vilkhu, V. Fan, M. Zaidi, C. Rhoades, N. Brackbill, G. Goetz, S. Wienbar, and the Stanford Artificial Retina team for helpful discussions. We thank R. Samarakoon and S. Kachiguine for technical assistance. We thank L. Jepson, P. Li, M. Greshner, G. Field, J. Gautier, A. Heitman, and G. Goetz for participating in data collection.

## Funding

NIH NEI F30-EY030776-03 (SM)

Polish National Science Centre grant DEC-2013/10/M/NZ4/00268 (PH)

Pew Charitable Trust Scholarship in the Biomedical Sciences (AS)

a donation from John Chen (AML)

Stanford Medicine Discovery Innovation Award (EJC)

Research to Prevent Blindness Stein Innovation Award (EJC)

Wu Tsai Neurosciences Institute Big Ideas (EJC)

NIH NEI R01-EY021271 (EJC)

NIH NEI R01-EY029247 (EJC)

NIH NEI P30-EY019005 (EJC)

